# Predicting Cortical Signatures of Consciousness using Dynamic Functional Connectivity Graph-Convolutional Neural Networks

**DOI:** 10.1101/2020.05.11.078535

**Authors:** Antoine Grigis, Chloé Gomez, Jordy Tasserie, Corentin Ambroise, Vincent Frouin, Béchir Jarraya, Lynn Uhrig

**Affiliations:** Université Paris-Saclay, CEA, NeuroSpin, 91191, Gif-sur-Yvette, France; Cognitive Neuroimaging Unit, Institut National de la Santé et de la Recherche Médicale U992, Gif-sur-Yvette, France; University of Versailles, Université Paris-Saclay, Neurosurgery Department, Foch Hospital, Suresnes, France

**Keywords:** deep learning, graph-convolutional network, self-supervised learning, cortical signatures of consciousness, resting-state fMRI, non-human primate

## Abstract

Decoding the levels of consciousness from cortical activity recording is a major challenge in neuroscience. Using clustering algorithms, we previously demonstrated that resting-state functional MRI (rsfMRI) data can be split into several clusters also called “brain states” corresponding to “functional configurations” of the brain. Here, we propose to use a selfsupervised machine learning method based on artificial neural networks to predict functional brain states across levels of consciousness from rsfMRI. The Functional Connectivity (FC) matrices reflect the brain-state dynamic at a given time. Because it is key to consider the FC topologies, a specific graph-Convolutional Neural Network (gCNN), namely BrainNetCNN, is considered to predict the brain states in awake and anesthetized nonhuman primates. To avoid the circularity that remains in the training stage, where the target is composed of pseudo-labels, recent self-supervised techniques are implemented. Using a linear probe for the prediction, the network achieves a prediction accuracy consistent with state-of-the-art methods lying in [0.655, 0.759] depending on the experimental settings. To put forward the interest of such a representation, the transition probabilities and the set of connections found to be important for predicting a brain state are computed. This latter is directly linked with the level of consciousness. The results demonstrate that deep learning methods are not only able to predict brain states but also provide additional insight into cortical signatures of consciousness with potential clinical consequences for the monitoring of anesthesia and the diagnosis of disorders of consciousness.

## 1 Introduction

A challenge in fundamental and clinical neuroscience is to predict the level of consciousness from brain activity recordings such as EEG or functional MRI (fMRI). Decoding levels of consciousness from the neural activity is essential for understanding the mechanism by which anesthetics induce consciousness loss, objectively monitoring the depth of anesthesia, and helping diagnose disorders of consciousness with higher accuracy [16]. Instead of a bottom-up investigation starting from molecules and cells, an information-processing analysis of brain functional imaging studies during the awake state or loss of consciousness induced by various anesthetics was proposed in non-human primates using dynamical resting-state [13, 8]. The resting-state fMRI (rsfMRI) data were divided into several clusters, also called “brain states”, corresponding to “functional configurations” of the brain and allowing decomposition of the consciousness conditions of a subject in terms of functional patterns. Briefly, a set of *k* Brain States {*BS*}^*k*^, corresponding to functional configurations, is obtained from nonhuman primates rsfMRI in the awake state and under different anesthetics. Then, each {*BS*}^*k*^ is ranked according to its similarity with the anatomical connectivity matrix defined in [4]. In [13, 8] the authors demonstrated that dynamic long-range functional correlation matrices reduce to the underlying structural anatomical connectivity matrix during anesthesia (Fig. 1). In other words, the brain states {*BS*}^*k*^ are directly related to the level of consciousness.

**Figure 1:**
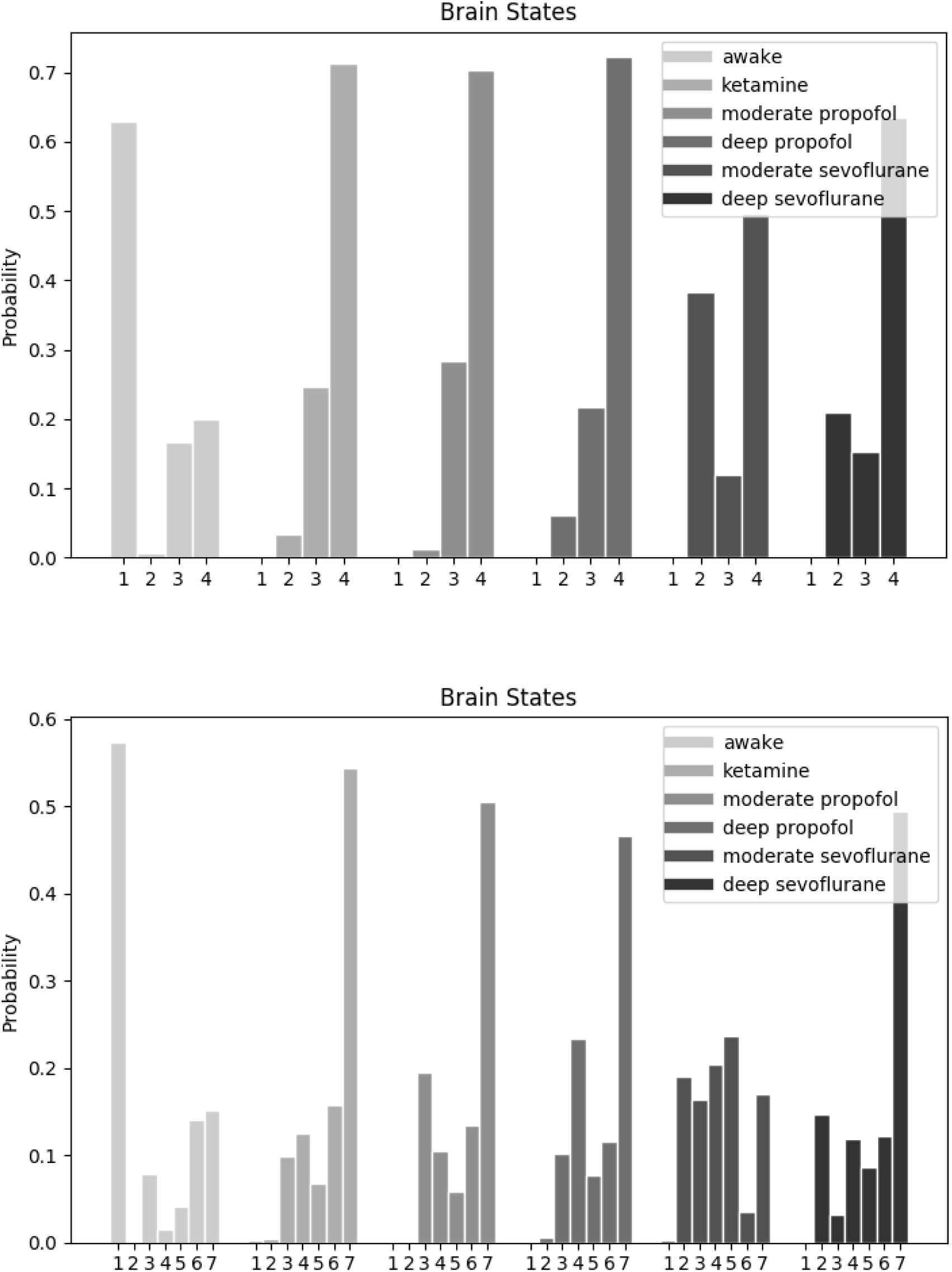
Probability distributions to observe in a rsfMRI time window, one of the ranked brain states {*BS*}^*k*^ for the awake and anesthesia conditions, and *k* = 4 or *k* = 7. Each bar represents the within-condition probability of occurrence of a state.

The methodology described above includes the entire dataset for the extraction and annotation of {*BS*}^*k*^. Although the set of {*BS*}^*k*^ is robust and stays stable when a new rsfMRI sequence is added to the database, the analysis process resumes the whole clustering, with potentially an appropriate initialization. Leveraging the increasing number of rsfMRI runs available, the objectives of this work are three-fold and will not require recomputing the {*BS*}^*k*^ set. First, we aim to confirm the interest of {*BS*}^*k*^ as labels to characterize the succession of states occurring under a given arousal condition. Using self-supervised machine learning methods, we will train a classifier that can predict one brain state from small rsfMRI temporal windows in completely new samples/runs [3]. In this work, we will use a specific graph-Convolutional Neural Network (gCNN) classifier named BrainNetCNN [11]. Standard CNNs can’t handle the spatial neighborhood of the Region Of Interests (ROIs) used to build the functional connectivity (FC) matrices. BrainNetCNN is composed of specific edge-to-edge, edge-to-node, and node-to-graph convolutional filters. These filters better reflect the topological locality of the ROIs used to generate the FC matrices. Another point in the training stage is to minimize the circularity between the targets composed of pseudo-labels and the predicted probabilities. For this purpose, recent contrastive learning techniques are implemented [20]. Second, we will challenge the data and learning process to model the brain dynamic oscillating from one state to another. Finally, we will leverage the embedded automatic differentiation of the deep learning framework to uncover which connections were used by the classifying network to be predictive of brain states. These predictive connections maps, which are duals of the {*BS*}^*k*^, yield some insights into both the brain states and the cortical signature of consciousness and can be considered (to some extent) as proxies of cortical signatures of consciousness.

## 2 Related works

### Unsupervised clustering

The objective of unsupervised clustering methods is to determine an internal grouping in a set of unlabeled data. In machine learning, many strategies can solve this task. Proposed methods can be grouped into four categories: 1) exclusive clustering, such as k-means, where each sample is assigned to one group, 2) overlapping clustering, such as fuzzy k-means, where each sample belongs to two or more clusters with different degrees of membership, 3) hierarchical clustering based on the fusion of nearest clusters starting from clusters containing only one sample, and 4) probabilistic clustering, such as mixtures of Gaussians, where each cluster is represented by a parametric distribution. These clustering machine learning techniques have been successfully applied in diverse applications such as the computation of the *BS*^*k*^ set [13].

### Deep self-supervised clustering

Recent works propose to uncover semantically relevant groups of samples using self-supervised deep learning techniques [12, 19]. Notably, Deep Cluster [12] alternates between a clustering phase and a back-propagation through a classification encoder driven by pseudo-labels. More specifically, at each epoch, previous clustering assignments produced by k-means on the generated embeddings are used as pseudo-labels for minimizing the cross-entropy. Such self-supervised techniques have proven to produce semantically valid clusters with no manual annotation. However, the training of these models is paved with pitfalls. The switch between the generation of pseudo-labels and the network training leads to clustering degeneracy and inconsistency. While the degeneracy issue is often solved with heuristics (which reassigns out-of-bounds samples/clusters), the inconsistency issue highly disrupts the training. For instance, in Deep Cluster [12], the clustering head is re-initialized at each epoch because k-means inevitably shuffles the assigned labels. In ODC [19], representations from a momentum encoder are stored, leading to a computationally costly nearest centroid problem at each batch. Such a training scheme is also sensitive to local minima because of the circular entanglement with the pseudo-labels.

### Deep contrastive learning

Another branch of the literature explores contrastive methods [14, 15, 20]. Distinctly, in SwAV [14], prototypes/clusters are discovered while enforcing consistency between cluster assignments of contrasted samples. Generic contrastive methods can also tackle the downstream clustering problem by applying a linear classifier on top of the fixed learned representation. For example, in SimCLR [15], input-distortion invariance is used as a pretext task to learn an appropriate representation of the data. Specifically, it encourages two augmented samples to be close in the representation space. An alternative option, computationally efficient and stable, has been proposed in Barlow Twins [20]. The learned representation is discovered by estimating the empirical cross-correlation matrix from contrasted samples. From a classification point of view, no constraint on class collision is enforced, thus allowing several samples sharing similar semantic content to be pulled apart. However, in practice, this problem is limited, and Barlow Twins produces clustering results as good or better than SwAV and SimCLR on ImageNet, Places-205, VOC07, and iNat18 [20].

## 3 Material and Methods

### 3.1 Material

This study uses the dataset described in [13]. Data were acquired either in the awake state or under different anesthetic conditions (ketamine, propofol, or sevoflurane anesthesia) in five macaque monkeys. Anesthesia levels were defined by a clinical score (the monkey sedation scale) and continuous electroencephalography monitoring. 156 rsfMRI runs with corresponding structural MRI were acquired on a Siemens 3-Tesla with a customized single transmit-receiver surface coil. The spatial prepossessing was performed with the NSM pipeline described in [18, 7], with specific time-series denoising operations [13, 8]. Specifically, voxel time series were filtered with a low-pass (0.05-Hz cutoff) and high-pass (0.0025-Hz cutoff) filters and a zero-phase fast-Fourier notch filter (0.03 Hz) to remove a pure artefactual frequency present in all the data. The variance of each time series was normalized to transform covariance into correlation matrices. Sliding windows are used to compute dynamic FC matrices reflecting the brain state at a given time [13]. Windowed segments of each time series were computed with a rectangular window (84 s width, which corresponds to 35 scans of 2.4 s each) and a sliding step of one scan, resulting in 464 windows per run. From the windowed rsfMRI, dynamic FC matrices were derived using the CoCoMac template composed of 82 cortical regions (41 cortical regions within each hemisphere) [13, 4]. The windowed rsfMRI time series from each region was converted into a connectivity matrix by projection into tangent space, capturing connectivity aspects from the correlation and partial correlation. The corresponding dynamic connectivity matrices led to a data array of shape 156 × 464 × 82 × 82 which was Fisher transformed to form a set of 156 × 464 samples. Each sample is more than a list of independent numbers and encodes the functional connectivity patterns. Dominant recurrent patterns of brain correlations (the {*BS*}^*k*^ set) were determined using an unsupervised k-means clustering with k states (or classes), k in [3, 9] as proposed in [13]. To avoid double-dipping (ie. the use of the same dataset for the training and the testing), the k-means was performed on a train set defined as a subset of the data. The labels associated with the remaining data were recovered by selecting the closest brain state {*BS*}^*k*^ forming the test set. To this end, the correlation coefficients between the test set dynamic FC matrices and each {*BS*}^*k*^ were computed, leading to a label array of shape 156 × 464 that will form the empirical ground-truth dataset.

### 3.2 Data partitioning

The dataset contains 5 monkeys with 156 rsfMRI runs in different arousal conditions (31 runs for the awake state, 25 runs for ketamine anesthesia, 25 runs for moderate propofol sedation, 30 runs for deep propofol anesthesia, 25 runs for moderate sevoflurane sedation, and 20 runs for deep sevoflurane anesthesia). We propose to leave one monkey out as the test set (35 runs) and the four others as the train set (121 runs). We further checked that both train and test sets have matching proportions of arousal conditions and brain state labels. The test set was set aside and not used during training. The train set was further split into validation and training folds using a 3-folds stratified cross-validation scheme.

### 3.3 Model architecture

The BrainNetCNN [11] works specifically with network data and will enforce the FC patterns. The model architecture is composed of convolutional layers followed by fully connected layers. Among the proposed configurations, we selected the E2Enet-sml, which consists in removing one edge-to-edge layer and two of the fully connected layers (Fig. 2). This configuration has shown excellent performances with a restricted number of parameters. Precisely, E2Enet-sml has an edge-to-edge layer composed of 32 1 × 82 and 32 82 × 1 filters producing feature maps of size 32 × 82 × 82, followed by an edge-to-node layer with 64 1 × 82 × 32 filters yielding feature maps of size 64 × 82 × 1, a node-to-graph layer with feature maps of size 1 × 1 × 30, and a fully connected layer with an output of size k. Increasing the number of feature maps with each layer is a common strategy for CNNs to compensate for the reductions along the other dimensions. Every layer uses very leaky rectified linear units as an activation function with a negative slope of 1/3.

**Figure 2:**
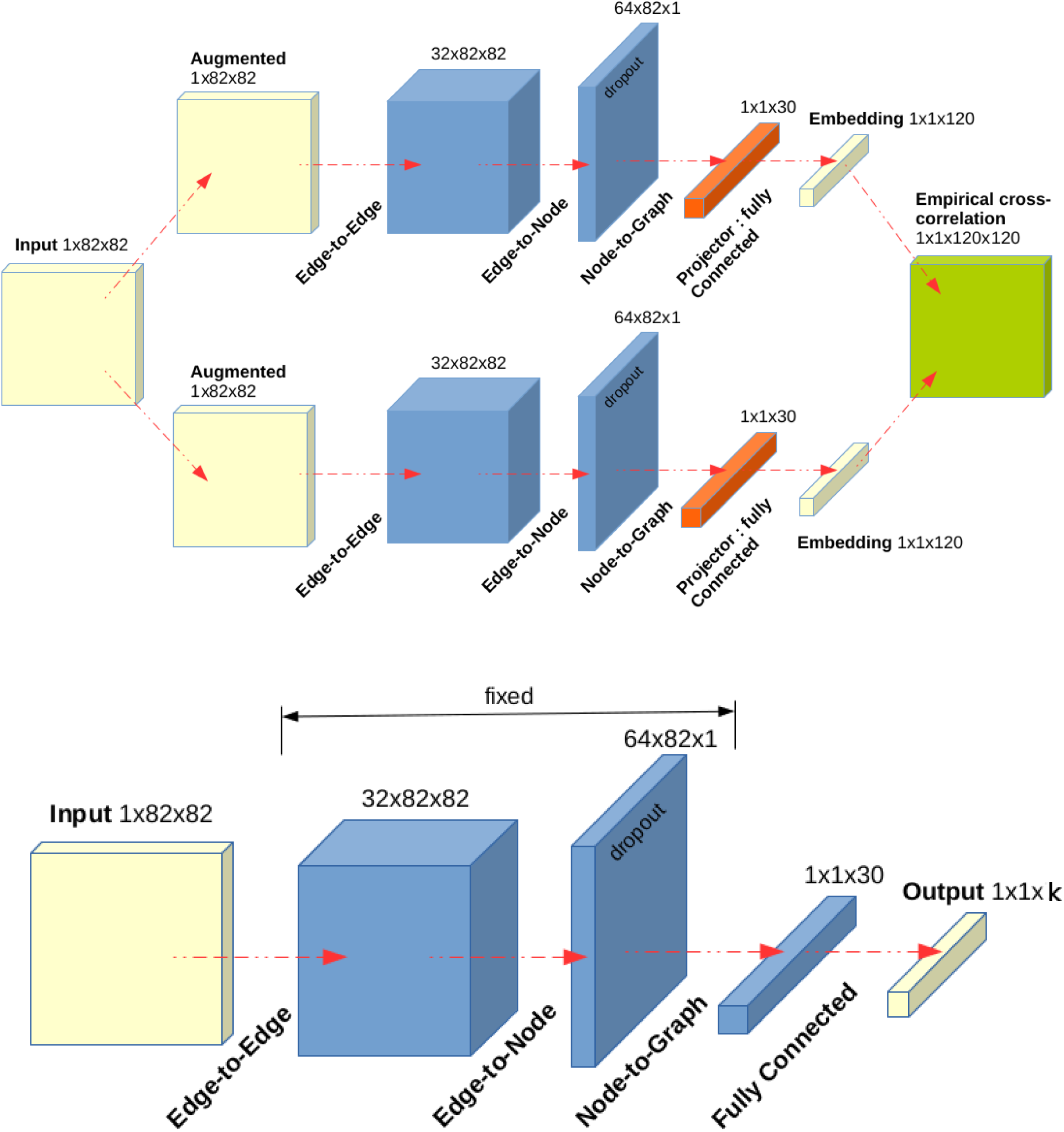
Schematic representation of the proposed training. The BrainNetCNN E2Enet-sml layers are represented in blue. In the first row the Barlow Twin contrastive approach is described. The fully connected layer of the BrainNetCNN is replaced by a projection head (in orange), and empirical cross-correlation matrix (in green) is computed from augmented/distorted samples (in yellow). In the second row the connected layer of the BrainNetCNN is trained using the k-means pseudo-labels by fixing the other weights.

Another objective is to minimize the impact of the pseudo-labels derived from the k-means during the training. This circularity issue is problematic, and without particular care, the weights of the CNN may more reflect the clustering algorithm than the variability of the data. To circumvent this limitation, we propose to use self-supervised contrastive learning techniques, more specifically, the Barlow Twins strategy [20]. The BrainNetCNN fully connected layer is replaced by a two hidden layers projection head leading to an embeddings space of size 120. The objective function tries to make the empirical cross-correlation matrix computed from twin embeddings (ie., outputs of the network fed with augmented/distorted versions of a sample) as close to the identity matrix as possible (Fig. 2). The FC matrices are distorted using random connections erasing scheme. The proportion of zeros is chosen randomly in the interval [0.3, 0.7]. The pseudo-labels from the k-means are now only used to train a linear classifier on the fixed representations learned by the Barlow Twins. All other weights are frozen to minimize the circularity between the targets and the predicted probabilities.

For training, we employed a dropout of 0.5 before the node-to-graph layer as shown in Fig. 2, and we follow the optimization protocol described in [20]. We use a LARS optimizer, a mini-batch of size 1024, a weight decay of 1e-6, a learning rate of 0.2 for the weights and 0.0048 for the biases and batch normalization parameters, and train the model for 1000 epochs. We use a learning rate warm-up period of 10 epochs, after which we reduce the learning rate by a factor of 1000 with a cosine decay schedule. The biases and batch normalization parameters are excluded from LARS adaptation and weight decay. The linear classifier is trained from the fixed representation, optimizing the cross-entropy loss during 100 epochs with an SGD optimizer, a weight decay of 1e-6, a learning rate of 0.01, and a cosine annealing schedule.

### 3.4 Maps of Predictive Connections

We use saliency maps to uncover the connections learned by the BrainNetCNN to be predictive of consciousness. Saliency maps are a popular visualization tool for gaining insight into why a deep learning model made an individual decision, such as classifying an image or a FC matrix [6, 10]. From a single backpropagation, the gradients of the target class w.r.t. the input FC matrix are generated from the first convolution layer. These gradients are used to visualize an FC-specific class saliency map, giving some intuition on connections within the input that contribute the most and least to the corresponding prediction. The badly predicted FC matrices are discarded, and resulting saliency maps are averaged brain-states-wise. In our application, they are helpful to extract proxies of the cortical signature of consciousness. These consciousness markers are rendered using a circular graph layout. The 41 cortical regions within each hemisphere are grouped, and the top *p* = 15% largest positive and negative connections are displayed. This threshold is computed at the brain-state level or across all brain states.

## 4 Results

### 4.1 Optimal number of the brain states

Before the presentation of the results on the three objectives of our work, we detail our sound estimation of the optimal number of brain states in the k-means. The basic idea behind the k-means consists in defining k clusters such that the within-cluster variations are minimum. Different metrics can be employed to select the optimal number of brain states among which we have retained the Elbow [5], the Silhouette [5, 2], and the Calinski and Harabasz [1] scores. The Elbow value is defined as the Within-Cluster Sum of Squared (WCSS) error for different values of k. The optimal k corresponds to the point of inflection on the WCSS versus k curve. Note that the Elbow score is more a decision rule than a metric. The Silhouette value measures how similar a sample is to its cluster compared to others and lies in [-1, 1]. The Calinski and Harabasz value is defined as the ratio between the within-cluster and the between-cluster dispersions. The last two scores are maximized with k. From the proposed experiment, we found no clear rule to select the optimal number of brain states for the k-means clustering (Fig. 3). The Silhouette and the Calinski and Harabasz indicators would select the smallest k value. However, the choice of k is the choice of the number of brain states that needs to be driven by biomarkers.

**Figure 3:**
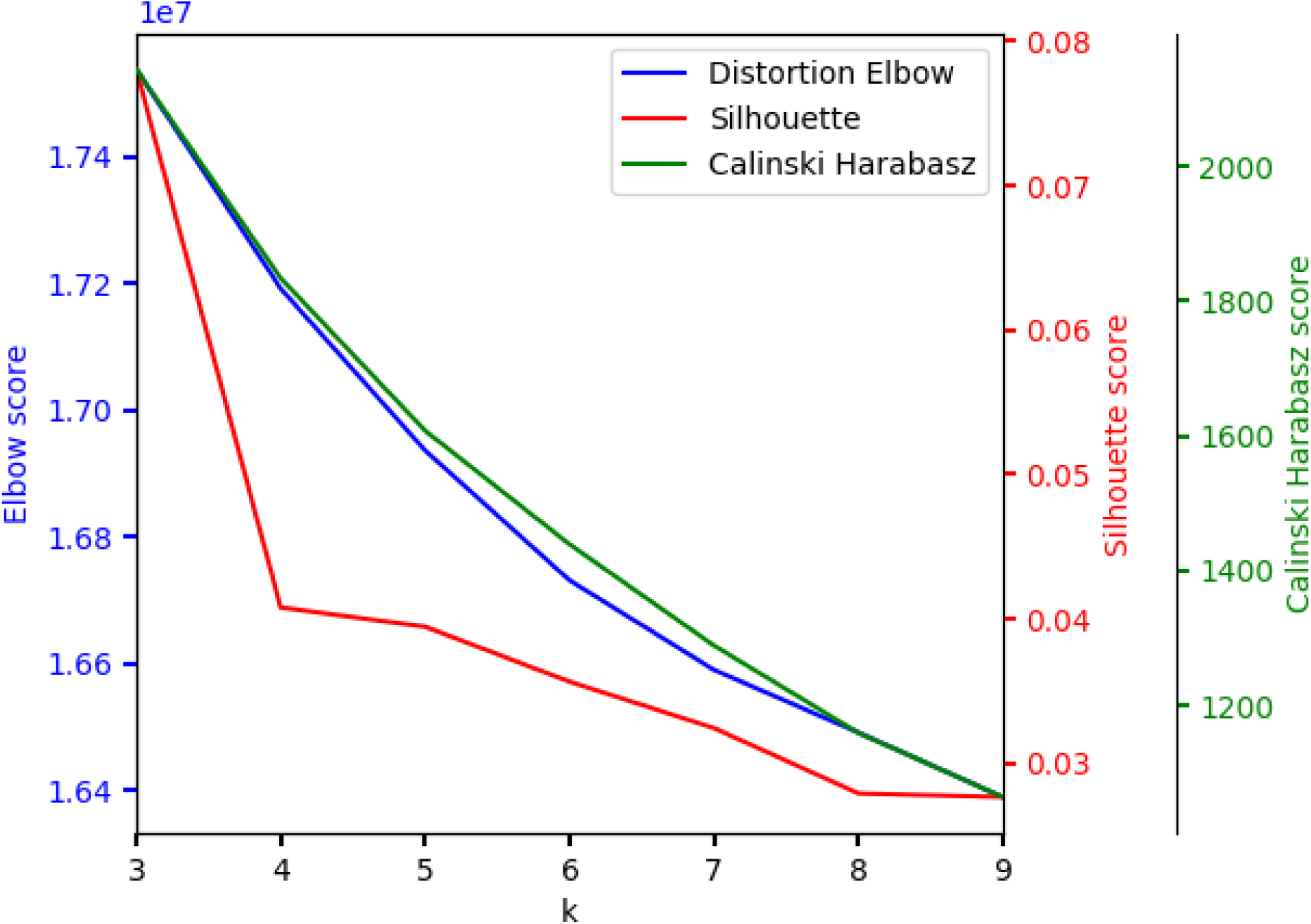
Determining the optimal number of brain states for the k-means clustering using the Elbow, the Silhouette, and the Calinski and Harabasz scores.

### 4.2 Performance of the classifier

Thus, we decided to monitor the BrainNetCNN prediction accuracy through different values of k (or targeted brain states {*BS*}^*k*^) as shown in Fig. 4. In a k-classes classification problem, the theoretical chance level is 1/k. As described in [9], this threshold holds for an infinite number of samples, and the smaller the sample size, the more likely chance performance differs from this theoretical chance level. In our application, the sample size is large (56,144 samples), and we assume that the chance level is close to its theoretical value. For all k values, we noticed that the accuracy is much higher than the theoretical chance level, demonstrating the interest of {*BS*}^*k*^ as labels to characterize the succession of states occurring under a given arousal condition. Smaller k values give better prediction results regardless of the chance level. The overall predictions are satisfactory and lie in [0.655, 0.759]. We put these results in perspective by training a machine learning linear SVC classifier directly on the input upper triangular FC data (Fig. 4). Both methods give similar results with a lower variability for the deep learning prediction when k increases. This result seems disappointing but reveals that the addressed downstream classification task is simple. Indeed, the annotations are generated using a k-means clustering, which consists of linear decision boundaries. While the proposed network uses a cascade of many layers of nonlinear processing units for feature extraction, the performance capabilities of the linear clustering task are likely limited by the nature of these empirical annotations. To go further, we rendered the machine learning and deep learning feature spaces using a Uniform Manifold Approximation and Projection (UMAP) [17]. The corresponding spaces have 3403 (the upper triangular elements of the FC matrices) and 30 (the latent space dimension of the BrainNetCNN) dimensions, respectively. We believe that the low dimensional features learned by the BrainNetCNN capture all the relevant information and will ease the interpretation as illustrated in Fig. 5.

**Figure 4:**
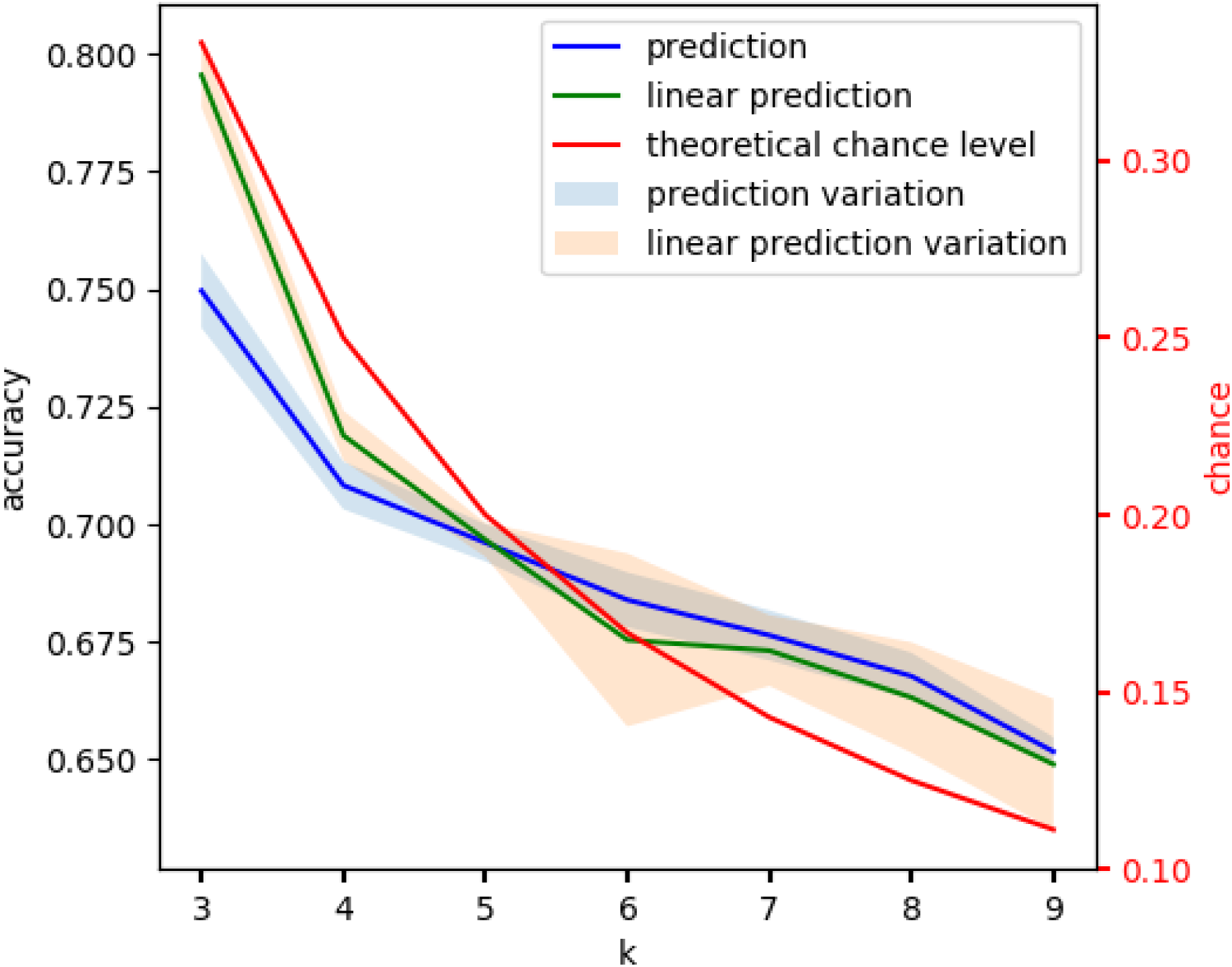
Determining the optimal number of brain states for the BrainNetCNN prediction using the accuracy as a reference metric: the mean and standard deviation computed across the training folds are displayed. A linear SVC prediction, as well as the theoretical chance levels are displayed as baselines. Note that results are always much higher than random chance.

**Figure 5:**
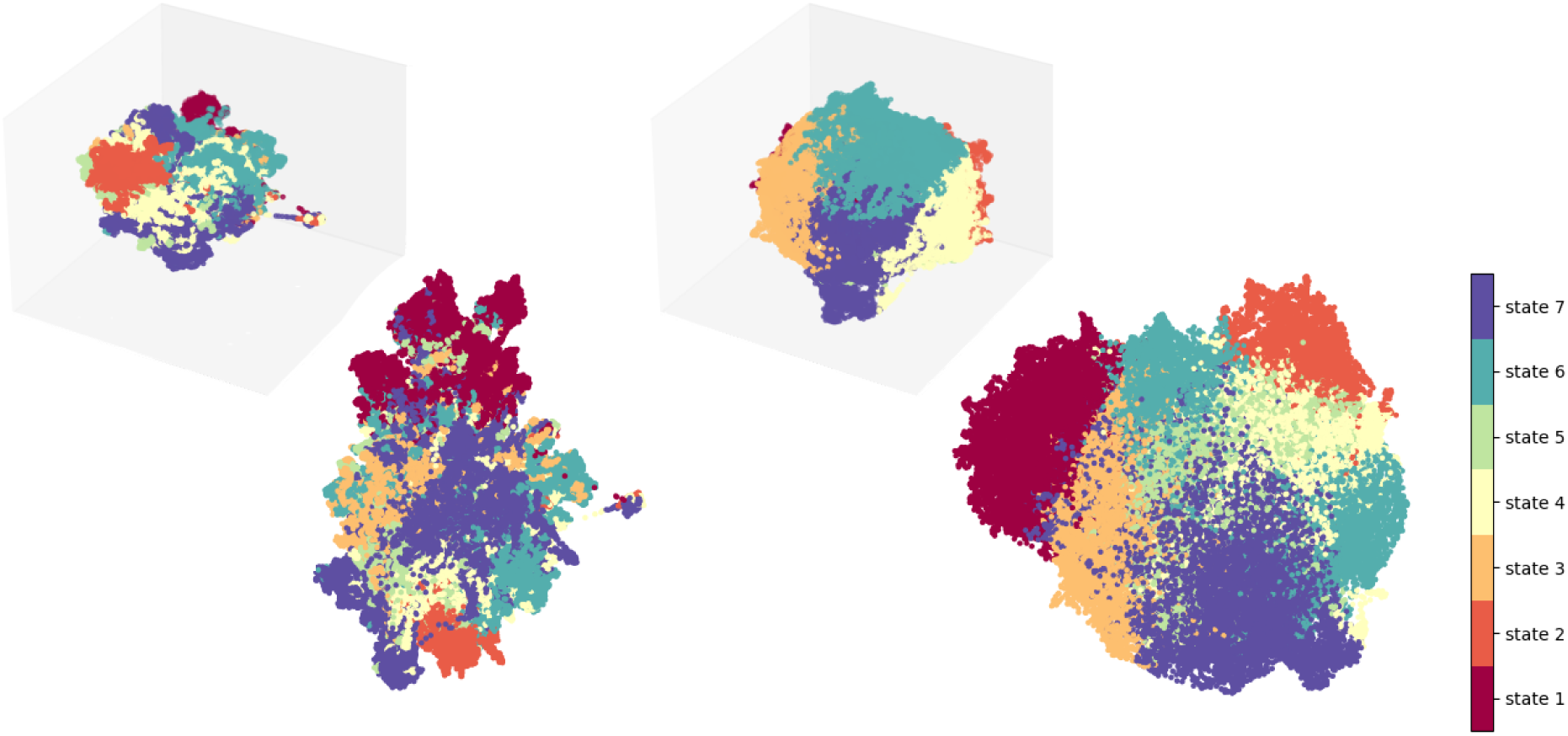
UMAP 2-d and 3-d projections of the machine learning (left) and deep learning (right) feature spaces: the upper triangular elements of the FC matrices (3403) vs the latent space dimension of the BrainNetCNN (30). The samples are colored using the k=7 brain states labels.

For the remaining part of the paper, we retained a biologically credible number of brain states, ie. k=4 or k=7 [13]. We chose accuracy as the metric to select the best-trained model. Then, the Receiver Operating Characteristic (ROC) metric is used to evaluate the BrainNetCNN classifier output quality. ROC curves show the true positive rate on the y-axis and the false positive rate on the x-axis. An Area Under the Curve (AUC) of 1 corresponds to the ideal case where the false positive rate is zero, and the true positive rate is one. Multi-label classification requires the binarization of the predicted brain state outputs. First, one ROC curve is drawn per label, followed by micro and macro-averaging. We thus demonstrate that the BrainNetCNN predicts the brain states with high reproducibility (*>* 0.92 AUC) as shown in Fig. 6.

**Figure 6:**
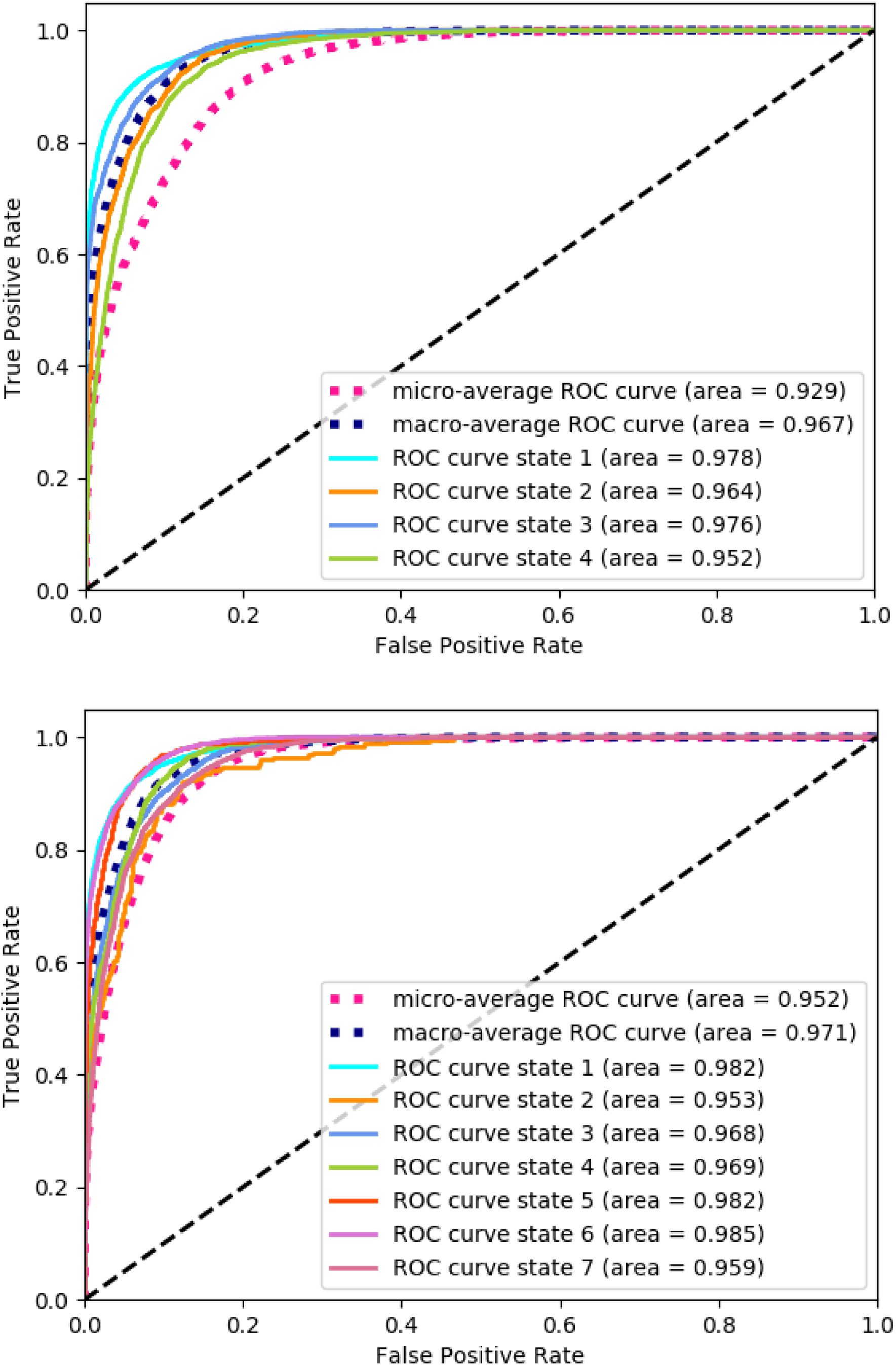
Evaluation of the BrainNetCNN classifier outputs quality using Receiver Operating Characteristic (ROC) curves for *k* = 4 and *k* = 7.

### 4.3 Towards modeling the brain states dynamic

Replaying the dynamic FC matrices movie for each run, the transitions probabilities to be in one state or another are directly accessible as the output of the trained network. Fig. 7 displays the predicted and the true labels as well as the network estimated probabilities for a specific run acquired in the awake condition. According to Fig. 1, in the awake condition, we expect a prevalence of state 1. The network decisions differ at state transitions. These chronograms give an insight into the monitored brain configurations, and in a prospective way model the brain dynamic oscillating from one state to another.

**Figure 7:**
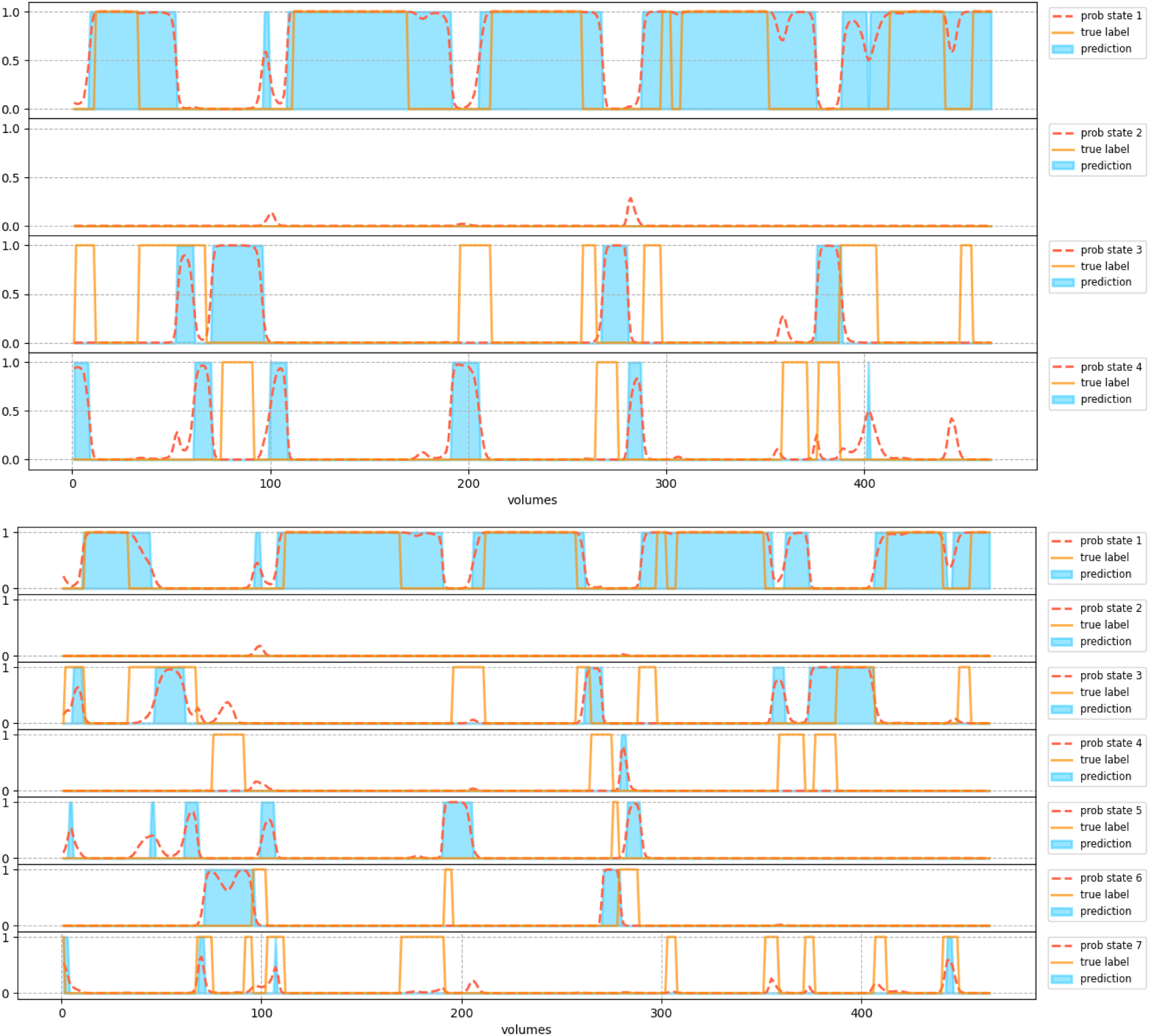
The BrainNetCNN predicted transitions with the associated probabilities for a test set run acquired in the awake condition for *k* = 4 and *k* = 7.

### 4.4 Maps of predictive connections

Proxies of the cortical signatures of consciousness are computed as described in section 3.4. The resulting mean saliency maps for each brain state are displayed in Fig. 8. The top *p* = 15% positive and negative connections using a specific or global threshold across brain states are presented in Fig. 9. The CoCoMac regions are grouped into seven locations comprising the cingulate, frontal, gustatory, insular, occipital, temporal, and parietal cortex. The brain states are characterized by the dynamical exploration of a rich, fluctuating set of functional configurations. These dynamical properties constitute a marker of consciousness. More specifically, the network uses shorter connections to identify a brain state far from the structural connectivity as depicted in Fig. 10. Let’s focus more closely on the two dominant brain states characterizing wakefulness and anesthesia, *BS*^1^ and *BS*^7^, respectively (cf. Fig. 1). The connections contributing the most to the corresponding predictions have an average Euclidean distance of 24.27 and 33.15, while the connections contributing the least have an average Euclidean distance of 30.55 and 21.05. This finding seems in line with the hypothesis formulated in [13] stating that dynamic FC matrices during anesthesia reduce to the underlying structural anatomical connectivity (ie. long-range connections), while during wakefulness closer auxiliary links are involved.

**Figure 8:**
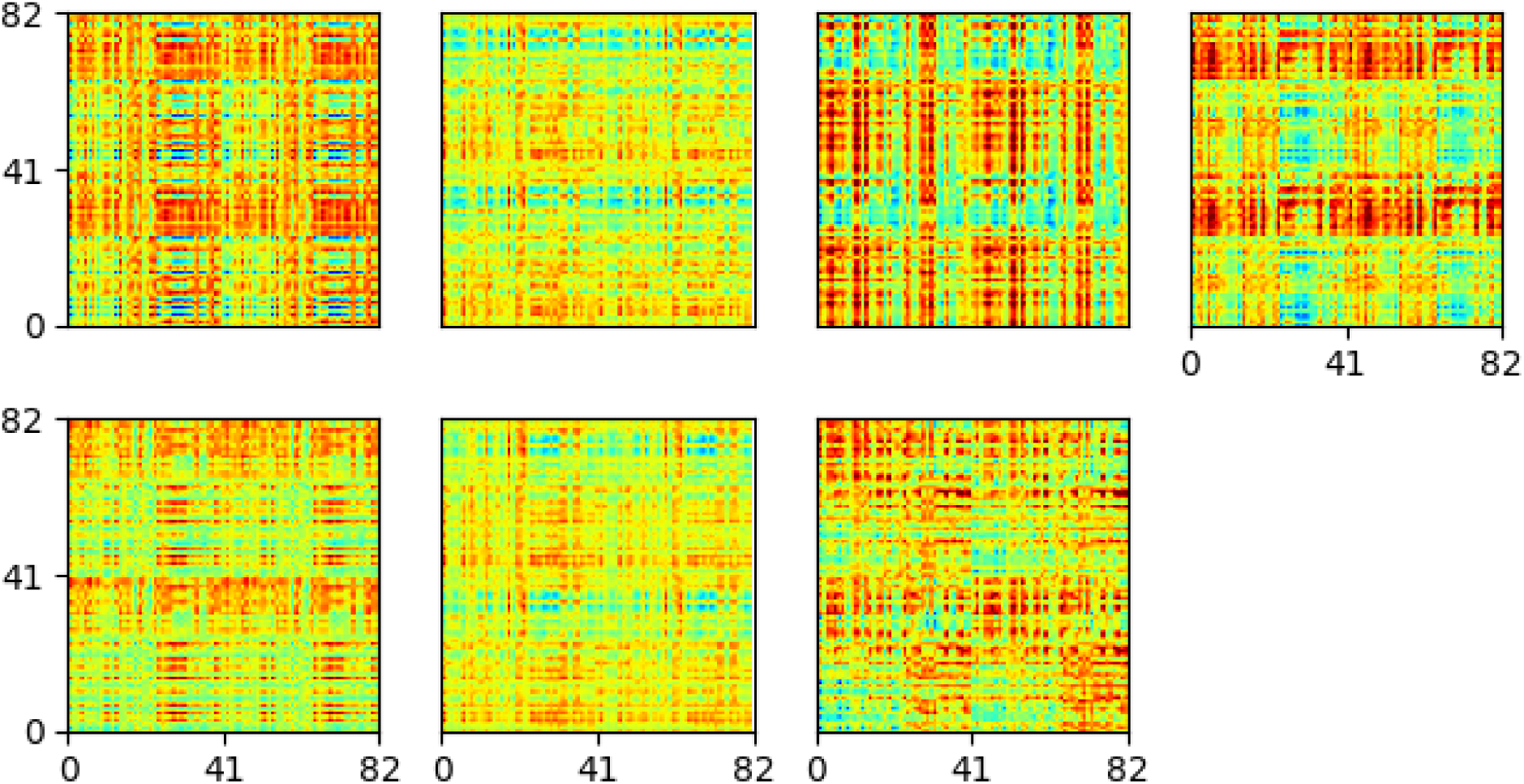
Resulting mean saliency maps for each brain state.

**Figure 9:**
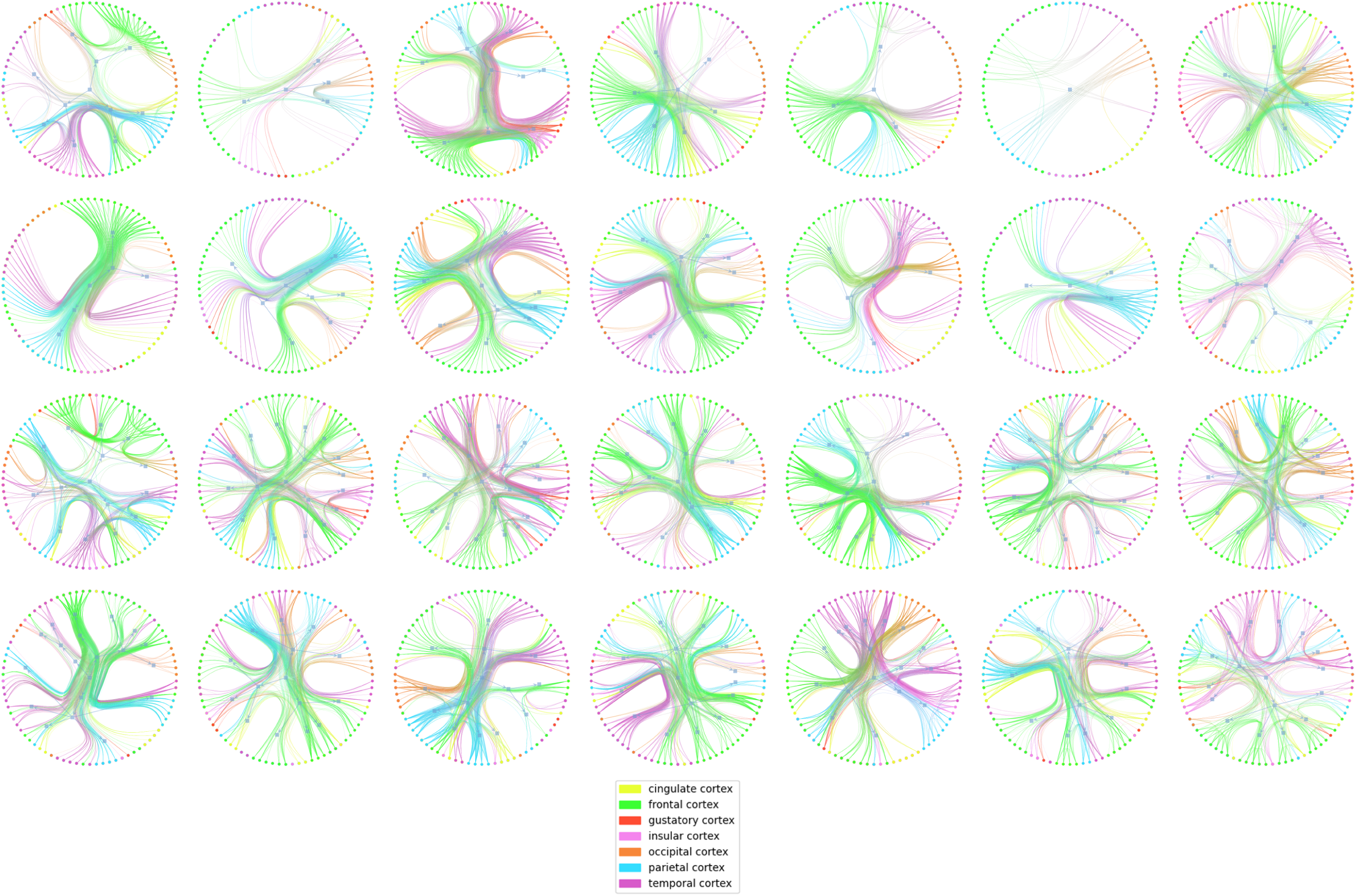
Within-brain states marker of consciousness for k=7, rendered using a circular graph where the top *p* = 15% positive and negative connections using a specific or global threshold across brain states are presented. Top to bottom rows contain the global-positive, global-negative, specific-positive, and specific-negative results, respectively.

**Figure 10:**
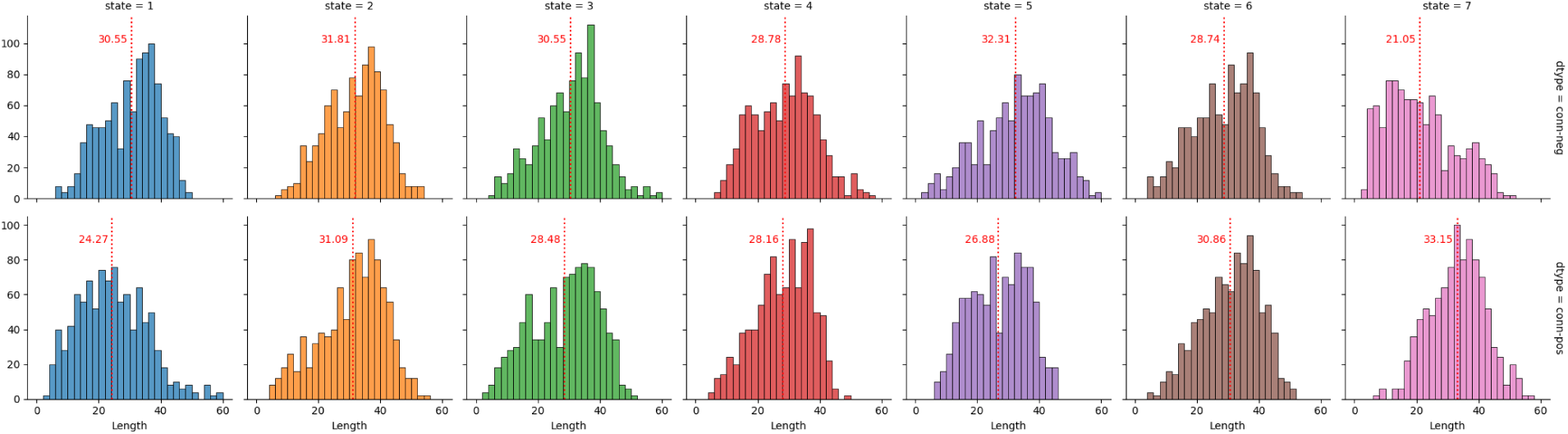
Within-brain states marker of consciousness for k=7, rendered using the top *p* = 15% positive and negative connections lengths (a global threshold is used). The means of each distribution are highlighted with dashed red lines. Top to bottom rows contain the positive, and negative connections of interest, respectively.

## 5 Discussion and conclusion

The BrainNetCNN gCNN model can predict brain states with good reproducibility and accuracy. It gives similar performance to a linear SVC on the input dynamical FC data pointing out the simplicity/restriction induced by the downstream classification task driven by the k-means pseudolabels. Nonetheless, the proposed network can scale to more complex tasks by learning complex representations. Using a self-supervised contrastive learning strategy reduces the circularity with the pseudo-labels. Nevertheless, this suited framework may suffer from a too coarse augmentation strategy. We also believe that the learned low-dimensional representation eases further interpretation. Interestingly, the BrainNetCNN predictions differ from the k-means ones at brain state transitions. More than a simple prediction tool, the proposed network can model the brain dynamic oscillating from one state to another and generate state signatures as sets of dominant connections.

As presented in [13], the entropy of the state transition matrix, the distance of each state to the underlying anatomical connectivity matrix, and the presence of negative correlations are helpful candidates when trying to find signatures of consciousness. We illustrate how the maps of predictive connections can disentangle which connections are useful to discriminate between different levels of wakefulness and bring significant insight into the brain states. These maps should help in understanding the brain states’ signature of consciousness. In future work, we want to apply this technique to anesthesia and disorders of consciousness clinical data.

In future work, we also want to systematically compare the performance of different classifiers based on machine learning or deep learning models to predict brain states. We believe that gCNN such as the BrainNetCNN will outperform models that do not model the topological locality of the regions used to generate the FC matrices. We also want to investigate the impact of enforcing consistency between cluster assignments on the learned representations as in SwAV [14].

